# A Molecular Pathway for Arterial-Specific Association of Vascular Smooth Muscle Cells

**DOI:** 10.1101/2019.12.27.889782

**Authors:** Amber N. Stratman, Margaret C. Burns, Olivia M. Farrelly, Andrew E. Davis, Wenling Li, Van N. Pham, Daniel Castranova, Joseph J. Yano, Lauren M. Goddard, Oliver Nguyen, Marina Venero Galanternik, Timothy J. Bolan, Mark L. Kahn, Yohsuke Mukouyama, Brant M. Weinstein

## Abstract

The preferential accumulation of vascular smooth muscle cells on arteries versus veins during early development is a well-described phenomenon, but the molecular pathways underlying this polarization are not well understood. During zebrafish embryogenesis the *cxcr4a* receptor (mammalian CXCR4) and its ligand *cxcl12b* (mammalian CXCL12) are both preferentially expressed on arteries at time points consistent with the arrival and differentiation of the first vascular smooth muscle cells (vSMCs). We show that autocrine *cxcl12b/cxcr4* activity leads to increased production of the vSMC chemoattractant ligand *pdgfb* by endothelial cells *in vitro* and increased expression of *pdgfb* by arteries *in vivo*. Additionally, we demonstrate that expression of the well-characterized blood flow-regulated transcription factor *klf2a* in primitive veins negatively regulates *cxcr4/cxcl12* and *pdgfb* expression, restricting vSMC recruitment to the arterial vasculature. Together, this signaling axis leads to the differential acquisition of smooth muscle cells at sites where *klf2a* expression is low and both *cxcr4a* and *pdgfb* are co-expressed, i.e. arteries during early development.

## INTRODUCTION

Endothelial cells (ECs) and mural cells are the main cellular constituents required for assembly of the arterial vascular wall. ECs form a single cell-thick, lumenized tube that is in direct contact with blood cells, immune cells, and hemodynamic forces. The endothelium is then surrounded by mural cells, including vascular smooth muscle cells (vSMCs) and pericytes—perivascular cell populations that promote long-term vessel stabilization and regulate vascular tone. vSMCs are largely associated with large caliber vessels, in particular arteries, while pericytes are associated with smaller vessels such as those in capillary beds. Arterial associated vSMCs provide tensile strength to the vascular wall by countering blood flow forces coming out of the heart, promoting maintenance and assembly of the vascular basement membrane, and helping regulate blood pressure (Bergers and Song, 2005; Hall-Glenn et al., 2012; Hedin et al., 1999; Heickendorff, 1989; Le et al., 2011; Lee et al., 2019; Leveen et al., 1994; Lilly, 2014; Lindahl et al., 1997; Osei-Owusu et al., 2014; Santoro et al., 2009; Scheppke et al., 2012; Stratman and Davis, 2012; Stratman et al., 2009; Stratman et al., 2017). Although vSMCs are critical modulators of arterial vasculature function, very little is known about the molecular cues that direct their preferential association with arteries rather than veins during early development. This preference has largely been thought to be mediated by differences in blood flow patterns, rates, and shear stresses associated with the arterial vasculature versus the venous vasculature (Ando et al., 2016; Chang et al., 2012; Lin and Lilly, 2014; Roostalu and Wong, 2018; Stratman et al., 2017). Although flow-mediated cues are likely critical, the cellular effectors mediating responses to blood flow and shear stress have yet to be fully elucidated.

The Klf2 transcription factor is one of the best-known blood flow-regulated genes, making it an attractive candidate for a role in flow-based regulation of vSMC recruitment. Klf2 is heavily expressed by postnatal endothelial cells and immune cells, and it has been studied extensively for its connection to atherosclerosis (Arkenbout et al., 2003; Atkins and Jain, 2007; Dekker et al., 2002; Dekker et al., 2005; Goddard et al., 2017; Huddleson et al., 2004; Kuo et al., 1997; Lee et al., 2006; Methe et al., 2007; Nayak et al., 2011; van Thienen et al., 2006). In atherosclerotic disease, sites of low Klf2 expression are prone to lesion development, hinting that suppressed Klf2 mediated signaling is linked to an activated state of cells comprising the vascular wall (Davies et al., 2013; Dekker et al., 2006; Dekker et al., 2005; Gimbrone and Garcia-Cardena, 2013; Goddard et al., 2017; Komaravolu et al., 2015; Mack et al., 2009; Parmar et al., 2006). Indeed, *in vitro* models suggest that EC induction of Klf2 could be a negative regulator of adjacent vSMC motility via promoted cellular differentiation (Dekker et al., 2006; Franzoni et al., 2019; Mack et al., 2009). Studies in mice, zebrafish, and *in vitro* have implicated Klf2 in angiogenesis, remodeling of the aortic outflow tract, heart formation and valve formation, and vascular stabilization during development (Adam et al., 2000; Bhattacharya et al., 2005; Chiplunkar et al., 2013; Dekker et al., 2002; Goddard et al., 2017; Kuo et al., 1997; Lee et al., 2006; Lin et al., 2010; Liu et al., 2005; Mack et al., 2009; Methe et al., 2007; Novodvorsky and Chico, 2014; Sangwung et al., 2017; Wu et al., 2008). However, the role of Klf2 in vSMC biology and function, and how it interfaces with known pathways and factors regulating vSMC recruitment and differentiation, remains largely unexplored.

Platelet Derived Growth Factor B (PDGFB) is one of the best-described factors regulating vSMC/mural cell biology. PDGFB ligand is produced by endothelial cells and it signals to mural cells via the PDGFRB receptor. A variety of studies have shown that this signaling pathway is critical for mural cell recruitment and proliferation, including the original reports demonstrating that PDGFRB knockout mice have decreased mural cell coverage of vessels and increased vessel dilation (Abramsson et al., 2007; Hellstrom et al., 1999; Leveen et al., 1994; Lindahl et al., 1997; Lindblom et al., 2003; Roostalu and Wong, 2018; Stratman and Davis, 2012; Stratman et al., 2017; Stratman et al., 2010). However, while recruitment of vSMCs is almost exclusively restricted to arteries during early development, PDGFB ligand is initially expressed on both primitive veins and arteries, before becoming largely restricted to arteries (Claxton et al., 2008; Hellstrom et al., 1999; Payne et al., 2018; Stanczuk et al., 2015), suggesting that other factors may help to suppress PDGFB expression in veins to direct vSMC recruitment to arteries.

The CXCR4 chemokine receptor and its associated ligand CXCL12 (aka SDF1a) are both expressed heavily on arteries or in tissue directly adjacent to arteries during early development (Busillo and Benovic, 2007; Bussmann et al., 2011; Cha and Weinstein, 2012; Corti et al., 2011; Fujita et al., 2011; Li et al., 2013). Chemokine signaling has been reported to have effects on vascular development, including during formation of blood vessels, formation of lymphatics, and potentially differentiation and recruitment of mural cells (Busillo and Benovic, 2007; Bussmann et al., 2011; Cha et al., 2012; Doring et al., 2014; Fujita et al., 2011; Gupta et al., 1998; Harrison et al., 2015; Li et al., 2013; Noels et al., 2014; Sebzda et al., 2008; Siekmann et al., 2009; Stratman et al., 2011). Knockout of CXCR4 in mice results in defective arterial patterning in the developing skin, in particular lack of alignment with nerves, with associated defects in mural cell coverage of the vasculature (the effects on large vessels were not analyzed in these studies) (Li et al., 2013). Other studies have suggested more direct effects of chemokine signaling in helping mural cells maintain their de-differentiated status, allowing for increased cellular motility, proliferation, and synthetic/matrix producing phenotype (Doring et al., 2014; Hamdan et al., 2011, 2014; Noels et al., 2014; Shi et al., 2014). CXCR4 and PDGFB are both thought to be blood flow responsive genes, suggesting potential links to Klf2 (Busillo and Benovic, 2007; Bussmann et al., 2011; Corti et al., 2011).

In this report we elucidate a molecular pathway promoting preferential association of vSMC with arterial blood vessels. We demonstrate that CXCL12/CXCR4 chemokine signaling in the arterial endothelium promotes PDGFB ligand expression, driving arterial vSMC recruitment. We also show that venous expression of KLF2 after the onset of blood flow negatively regulates CXCR4 and PDGFB production to inhibit venous recruitment of vSMC. Together, our findings highlight a molecular pathway driving arterial-specific recruitment of vSMCs, via regulated and coupled expression of chemokines and PDGFB in the vasculature.

## RESULTS

### Smooth muscle cells preferentially associate with developing arteries

The presence of a thickened vascular wall with abundant vascular smooth muscle cells (vSMC) is perhaps the most clearly evident morphological feature distinguishing arteries from veins, but the molecular mechanisms responsible for preferential acquisition of vSMC by arteries remains largely unexplored. As we and others have previously reported, vSMC emerge from the medial sclerotome of the zebrafish trunk beginning at approximately 3 dpf, taking up residence around the closely juxtaposed dorsal aorta, but not around the equally closely adjacent cardinal vein (Fig. 1A;(Ando et al., 2016; Santoro et al., 2009; Stratman et al., 2017; Whitesell et al., 2014)). This differential recruitment of vSMC to the trunk dorsal aorta but not the cardinal vein is readily observed in 5 dpf *Tg(tagln:eGFP),Tg(kdrl:mCherry-CAAX)* double-transgenic zebrafish with mCherry-positive vascular endothelium (magenta) and eGFP-positive vSMC (green) (Fig. 1B,C). To identify candidate factors that might be playing a role in guiding the dorsal aorta-restricted recruitment of vSMC, we used the ZFIN gene expression database (zfin.org) to carry out an *in silico* survey for secreted factors differentially expressed in the dorsal aorta but not in the cardinal vein at approximately 2.5 dpf. The *cxcl12b* chemokine ligand (also known as SDF1a) and its receptor *cxcr4a* were identified as promising candidates. Although, expressed in both cardinal vein and dorsal aorta at earlier stages of development (1.25 dpf, data not shown), by 2.5 dpf when vSMC recruitment is beginning, *cxcl12b* and *cxcr4a* expression is strongly enriched in the dorsal aorta compared to the cardinal vein (Fig. 1D-G).

**Figure 1.**
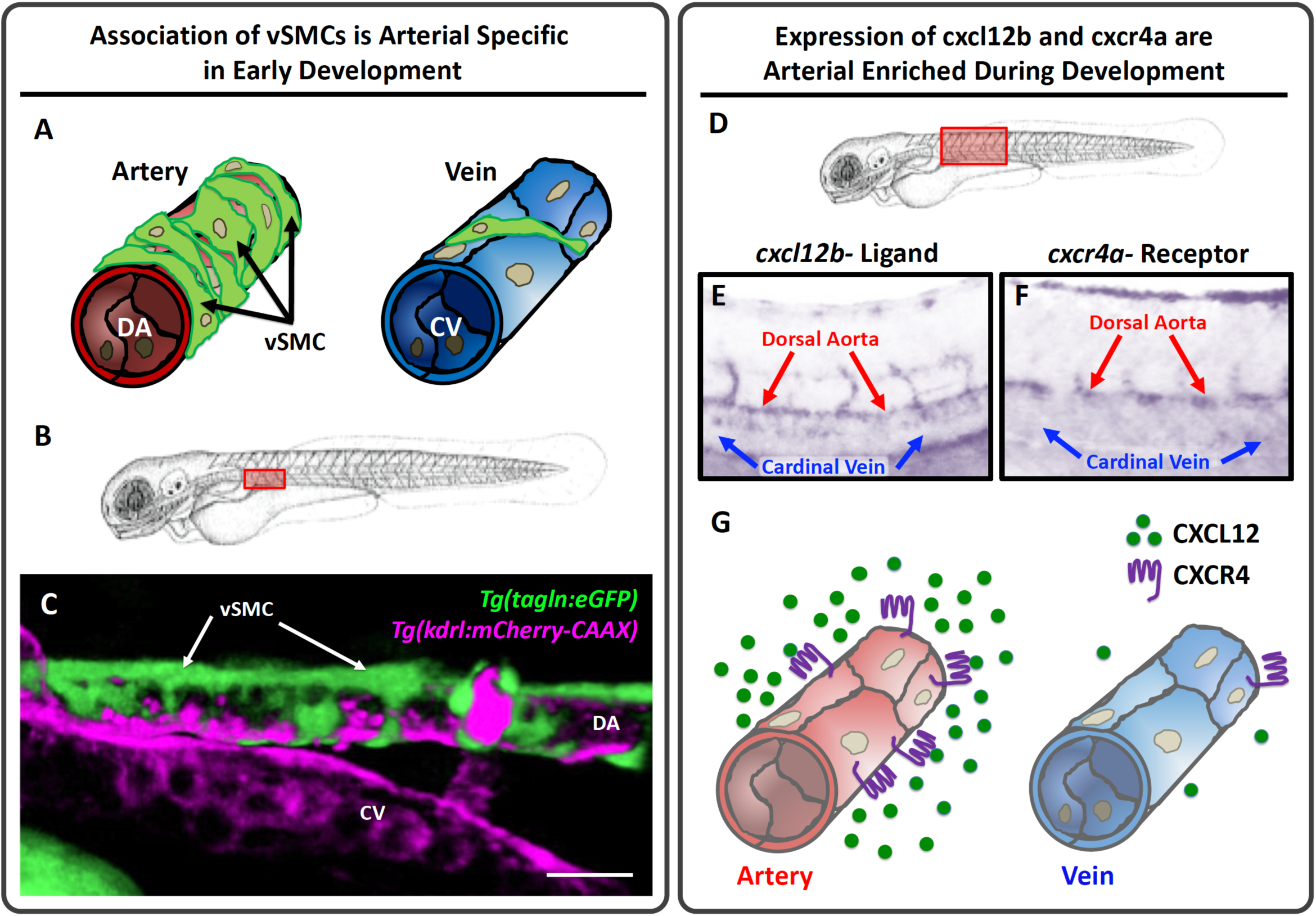
vSMCs associate with arteries during development. **A**, Schematic diagram illustrating vSMC coverage of arteries and lack of coverage of veins. **B**, Schematic diagram of a zebrafish larva with the red box highlighting the area imaged in panel C. **C**, Confocal micrograph of the anterior trunk of a 5 dpf *Tg(tagln:eGFP), Tg(kdrl:mCherry-CAAX)* double-transgenic zebrafish larva expressing eGFP in vSMCs (green) and mCherry-CAAX in the endothelium (magenta). vSMCs are associated with the dorsal aorta (DA) and not the cardinal vein (CV). **D,** Schematic diagram of a zebrafish larva with the red box highlighting the area imaged in panels E and F. **E,F**, Whole mount *in situ* hybridization of the mid-trunk of 48 hpf zebrafish larvae probed for *cxcl2b* ligand (E) or *cxcr4a* receptor (F). Expression of both genes is enriched in the dorsal aorta compared to the cardinal vein. **G**, Schematic representation of arterial-enriched expression of *cxcl12b* and *cxcr4a*. Scale bars = 75 µm (panel C).

### Chemokine signaling regulates vSMC association with arteries

To examine whether *cxcl12b/cxcr4a* chemokine signaling plays a role in dorsal aorta-specific vSMC recruitment, we used CRISPR/Cas9 technology to generate 24 bp and 7 bp deletion mutants in *cxcl12b* and *cxcr4a,* respectively (Supp. Fig. 1A,B). Homozygous *cxcl12b^Δ24/Δ24^* mutant animals show reduced numbers of vSMC associated with the dorsal aorta at 5 dpf (Fig. 2A-C) and an increased dorsal aorta diameter (Fig. 2D), a previously-reported consequence of defects in vSMC coverage of vessels (Stratman et al., 2009; Stratman et al., 2017). Homozygous *cxcr4a^Δ7/Δ7^* mutant animals also have reduced numbers of dorsal aorta-associated vSMC and increased dorsal aorta diameter (Fig. 2E-H). Deletion of the Cxcr4 gene in mice similarly results in decreased thickness of the dorsal aorta vSMC (smooth muscle actin-positive) wall (Fig. 2I-K) and dorsal aorta luminal enlargement (Fig. 2L) at E12.5, suggesting the role of chemokine signaling in vSMC recruitment is evolutionarily conserved. If chemokine signaling is indeed promoting vSMC recruitment to the dorsal aorta, we reasoned that forced mis-expression of *cxcl12b* in the cardinal vein (where it is not normally expressed at 3 dpf) might result in ectopic targeting of vSMC to this vessel (Fig. 3A,B). To test this, we injected *Tg(tagln:eGFP)* transgenic zebrafish with a Tol2 transgene containing the *mrc1a* promoter driving expression of *cxcl12b* and mCherry, co-translationally linked together via the 2A peptide sequence (Fig. 3A,B). We have previously shown that the *mrc1a* promoter drives robust expression in the early cardinal vein in addition to other vessels (Jung et al., 2017). Mosaic expression of *cxcl12b* in the cardinal vein (marked by mCherry expression) promoted ectopic cardinal vein recruitment of vSMC without affecting vSMC acquisition by the dorsal aorta (Fig. 3C-E). Together these gain- and loss-of-function experiments support the idea that dorsal aorta-restricted chemokine signaling is important for differential arterial recruitment of vSMC.

**Figure 2.**
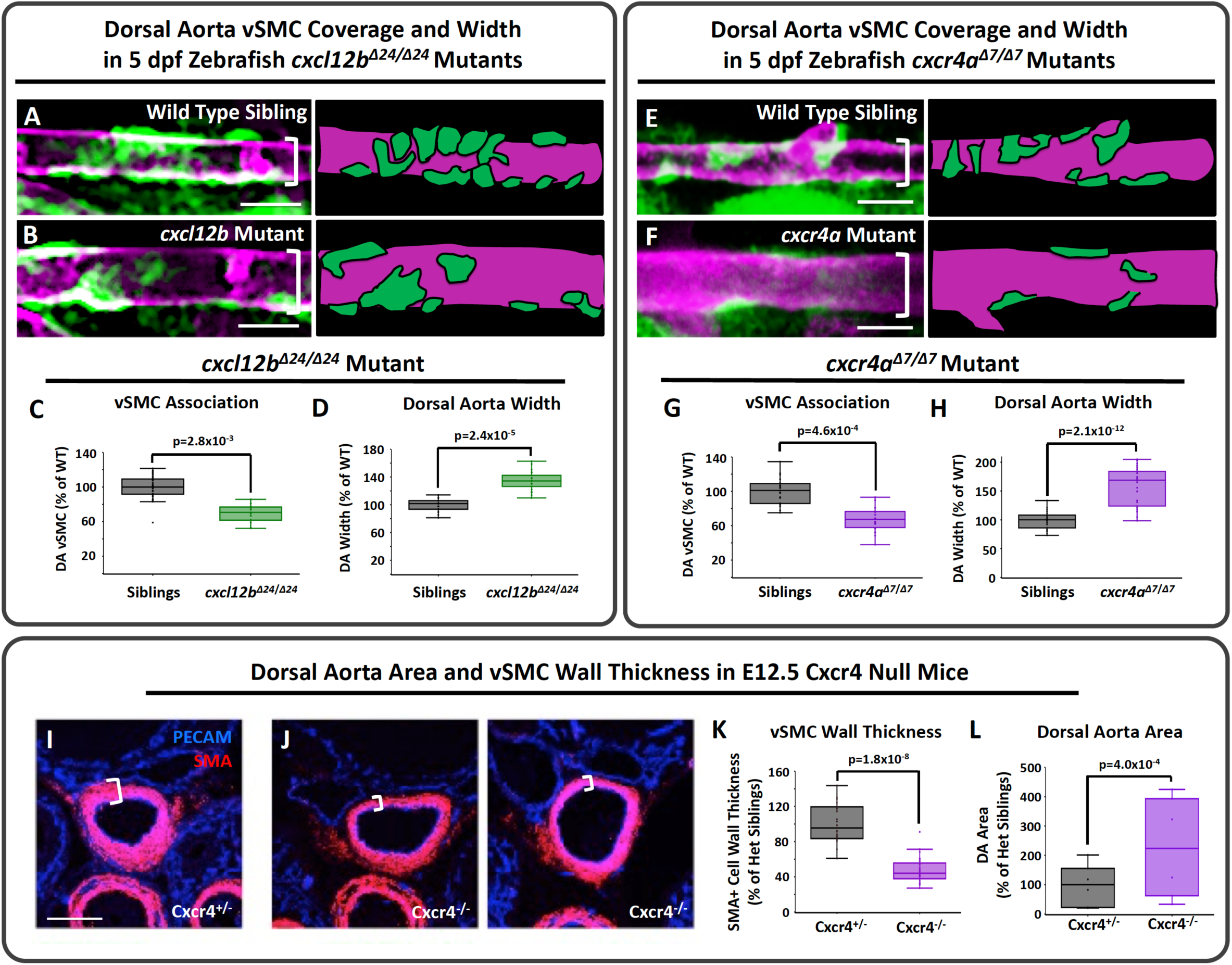
Disrupting *cxcl12b/cxcr4a* signaling decreases vSMC association with arteries. **A,B,** Confocal images (left) and schematic representations (right) of the dorsal aorta (DA) in the anterior trunk of 5 dpf *Tg(tagln:eGFP), Tg(kdrl:mCherry-CAAX)* double-transgenic sibling (A) or *cxcl12b^Δ24/Δ24^* mutant (B) zebrafish expressing eGFP in vSMCs (green) and mCherry-CAAX in the endothelium (magenta). **C, D,** Quantification of the number of associated vSMC (C) and width (D) of the dorsal aorta in 5 dpf *Tg(tagln:eGFP), Tg(kdrl:mCherry-CAAX)* double-transgenic sibling (black columns) or *cxcl12b ^Δ24/Δ24^* mutant (green columns) larvae. Values are expressed as a percentage of control siblings and averaged from three individual experiments. **E,F,** Confocal images (left) and schematic representations (right) of the dorsal aorta (DA) in the anterior trunk of 5 dpf *Tg(tagln:eGFP), Tg(kdrl:mCherry-CAAX)* double-transgenic siblings (E) or *cxcr4^Δ7/Δ7^* mutant (F) zebrafish expressing eGFP in vSMCs (green) and mCherry-CAAX in the endothelium (magenta). **G,H,** Quantification of the number of associated vSMC (G) and width (H) of the dorsal aorta in 5 dpf *Tg(tagln:eGFP), Tg(kdrl:mCherry-CAAX)* double-transgenic sibling (black columns) or *cxcr4^Δ7/Δ7^* mutant (purple columns) larvae. Values are expressed as a percentage of control siblings and averaged from three individual experiments. **I,J**, Confocal images of immunohistochemically stained transverse sections through the dorsal aorta of E12.5 Cxcr4+/- heterozygous sibling (I) and Cxcr4-/- mutant (J) mice, probed for platelet endothelial cell adhesion molecule-1 (PECAM) for endothelium (blue) and alpha smooth muscle actin (SMA) for vascular smooth muscle (vSMC, red). White brackets note the thickness of the vSMC layer surrounding the DA. **K,L,** Quantification of vSMC wall thickness (K) and lumenal area (L) of the dorsal aorta of E12.5 Cxcr4+/- heterozygous sibling (black columns) and Cxcr4-/- mutant (purple columns) mice, measured from immunohistochemically stained sections as in panels I and J. Values are expressed as a percentage of heterozygous siblings and averaged from three individual experiments. Scale bars = 75 µm (panels A,B,E,F), 100 µm (panels I,J). Box plots are graphed showing the median versus the first and third quartiles of the data (the middle, top, and bottom lines of the box respectively). The whiskers demonstrate the spread of data within 1.5x above and below the interquartile range. All data points are shown as individual dots, with outliers shown above or below the whiskers. P-values are indicated above statistically significant datasets.

**Figure 3.**
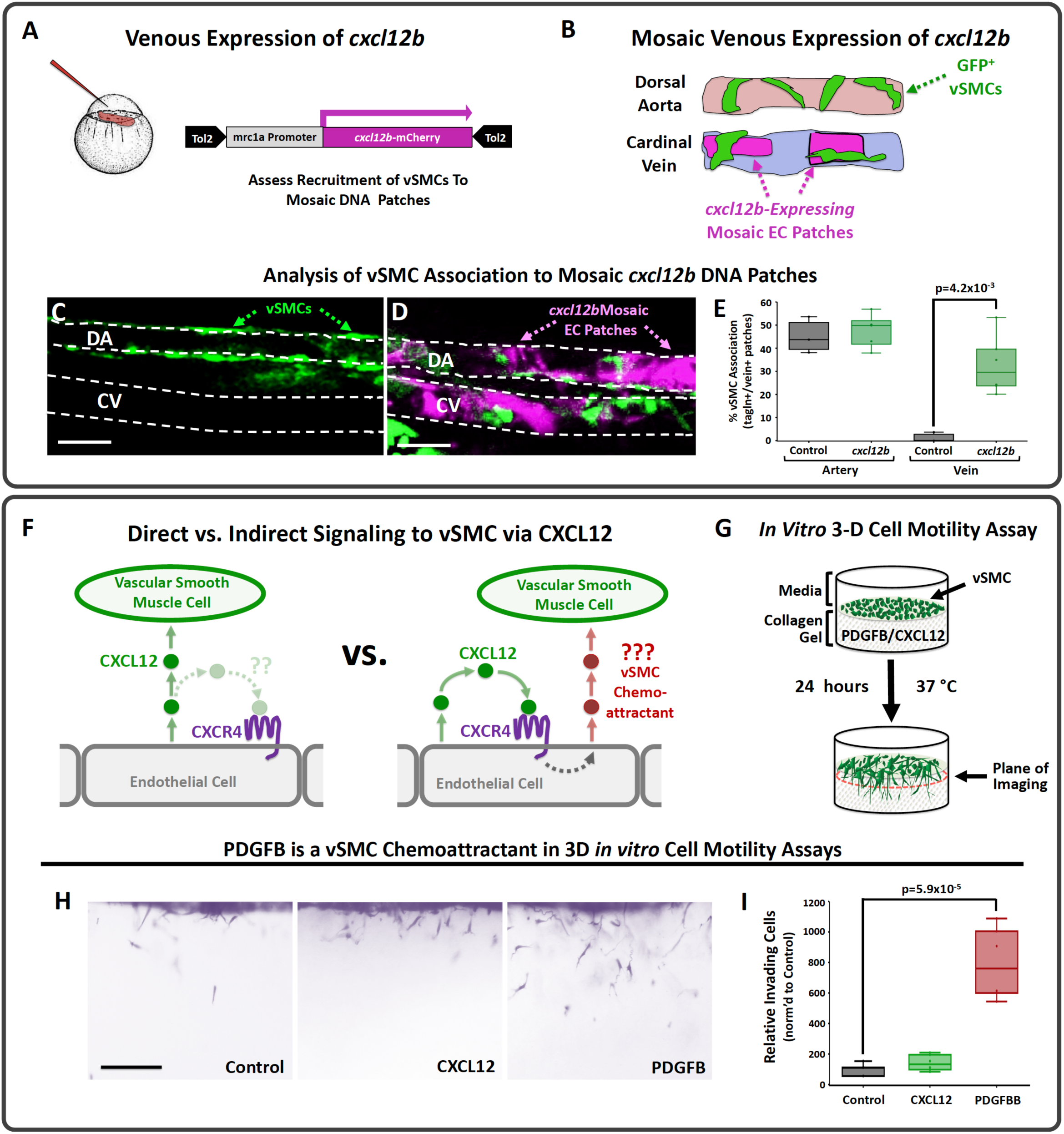
*cxcl12b* promotes vSMC association without serving as a direct chemoattractant. **A,B,** Schematic diagrams illustrating the experimental design for using the *mrc1a* promoter to drive ectopic mosaic expression of *cxcl12b* in veins. **A,** A *Tol2(mrc1a:cxcl12b-2a-mCherry)* DNA construct co-translationally expressing *cxcl12b* and mCherry under the control of the *mrc1a* promoter is injected into *Tg(tagln:eGFP)* transgenic zebrafish embryos at the 1 cell stage. **B,** At 4 dpf *tol2(mrc1a:cxcl12b-2a-mCherry)*-injected zebrafish larvae are analyzed for vSMC (eGFP) association at sites of mCherry (i.e. *cxcl12b*) expression in the dorsal aorta and cardinal vein. **C,D**, Representative confocal images of the mid-trunk of 4 dpf *Tg(tagln:eGFP)* transgenic larvae injected with either control *Tol2(mrc1a)* “empty vector” (C) or *Tol2(mrc1a:cxcl12b-2a-mCherry)* (D). eGFP-expressing vSMCs are shown in green, *cxcl12b-2a-mCherry* expression in dorsal aorta (DA) or cardinal vein (CV) endothelium is shown in magenta. **E**, Quantification of eGFP-positive vSMC associated with the dorsal aorta (DA) or cardinal vein (CV) in 4 dpf *Tg(tagln:eGFP)* transgenic zebrafish injected with either control *Tol2(mrc1a)* “empty vector” (black columns) or *Tol2(mrc1a:cxcl12b-2a-mCherry)* (green columns), showing strongly increased association of vSMCs with the cardinal vein. **F**, Schematic diagrams showing potential models for direct (left) versus indirect (right) mechanisms for promoting arterial recruitment of vSMC via CXCL12. **G**, Schematic diagram illustrating the 3D pulmonary artery smooth muscle cell (PASMC) motility assay. CXCL12, PDGFB, or nothing (control) is placed within the collagen gel to determine if PASMCs migrate towards these potential chemoattractants. **H,** Representative lateral images of 3D collagen gels showing PASMCs within the collagen matrix for each gel condition. **I**, Quantification of the relative number of PASMCs invading the collagen gel. The control is set to 100% and the CXCL12 and PDGFB conditions normalized to this level of invasion. Scale bars = 75 µm (panels C,D), 200 µm (panel H). Box plots are graphed showing the median versus the first and third quartiles of the data (the middle, top, and bottom lines of the box respectively). The whiskers demonstrate the spread of data within 1.5x above and below the interquartile range. All data points are shown as individual dots, with outliers shown above or below the whiskers. P-values are indicated above statistically significant datasets.

### PDGFB modulates vSMC association with the dorsal aorta

We carried out additional experiments to examine whether CXCL12 acts directly as a paracrine chemoattractant for arterial vSMC or indirectly via autocrine activation of arterial endothelial CXCR4 to induce expression of some other chemoattractant factor (Fig. 3F). Using a previously described *in vitro* 3-D cell motility assay (Fig. 3G; (Koh et al., 2008; Stratman et al., 2011; Stratman et al., 2009)), we showed that human coronary artery smooth muscle cells (CASMCs) have little or no chemoattractant activity for CXCL12, unlike PDGFB which has a robust activity in this assay (Fig. 3H,I). PDGFB is a well-documented *in vivo* vSMC chemoattractant expressed by endothelial cells and required for vSMC recruitment in mice (Abramsson et al., 2007; Hellstrom et al., 1999; Leveen et al., 1994). Zebrafish *pdgfb* is expressed weakly in both the dorsal aorta and cardinal vein at 1.5 dpf (Fig. 4A-C), but by the time vSMC recruitment begins at 3 dpf it shows preferential expression in the developing dorsal aorta (Fig. 4C). To examine the role of *pdgfb* in vSMC recruitment in the zebrafish we used CRISPR/Cas9 mutagenesis to generate mutants in the *pdgfbb* and closely related *pdgfba* genes (Supp. Fig. 1C). Zebrafish homozygous mutants for either *pdgfbb* or *pdgfba* alone display modest reductions in vSMC recruitment to the dorsal aorta at 5 dpf, but animals homozygous for the mutations in both genes display a strong reduction in vSMC recruitment (Fig. 4D-F). Together with previous findings from our lab using expression of dominant-negative *pdgfrb*-DN to disrupt vSMC recruitment (Stratman et al., 2017), these results suggest that, as noted in mice, PDGFB signaling serves as an important vSMC chemoattractant in the zebrafish.

**Figure 4.**
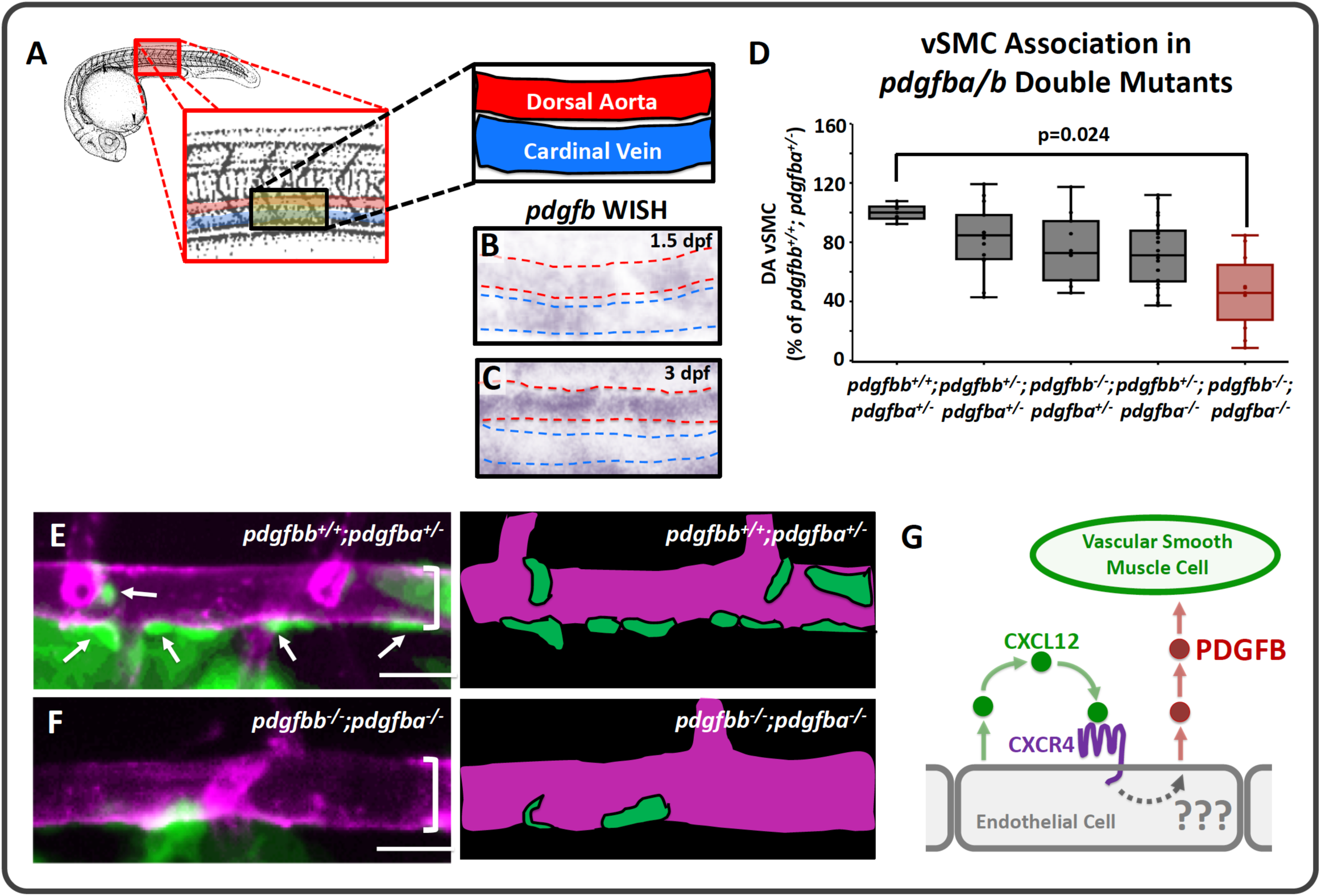
*pdgfb* mediated signaling regulates vSMC association with arteries. **A**, Schematic representation of the area imaged for *in situ* hybridization analysis, demonstrating the location of the dorsal aorta (DA) and cardinal vein (CV) within these images. **B,C**, Whole mount *in situ* hybridization (WISH) images of *pdgfb* transcript (purple) in 1.5 dpf (B) and 3 dpf (C) zebrafish. Red dashed lines outline the dorsal aorta, blue dashed lines outline the cardinal vein. **D**, Quantification of the number of vSMC associated with the dorsal aorta in 5 dpf *Tg(tagln:eGFP), Tg(kdrl:mCherry-CAAX)* double-transgenic zebrafish carrying different combinations of heterozygous or homozygous *pdgfba* and *pdgfbb* mutants. Values are averaged from three individual experiments and expressed as a percentage of the *pdgfba*+/-, *pdgfbb+/+* control. **E,F**, Confocal images (left) and schematic representations (right) of the dorsal aorta (DA) in the anterior trunk of 5 dpf *Tg(tagln:eGFP), Tg(kdrl:mCherry-CAAX)* double-transgenic *pdgfba*+/-, *pdgfbb+/+* sibling (E) or *pdgfba*-/-, *pdgfbb-/-* double homozygous mutant (F) zebrafish expressing eGFP in vSMCs (green) and mCherry-CAAX in the endothelium (magenta). **G**, Schematic diagram illustrating the proposed model for endothelial-autonomous chemokine signaling driving increased endothelial PDGFB ligand production, and thereby indirectly promoting vSMC acquisition by arteries. Scale bars = 75 µm (panels E,F). Box plots are graphed showing the median versus the first and third quartiles of the data (the middle, top, and bottom lines of the box respectively). The whiskers demonstrate the spread of data within 1.5x above and below the interquartile range. All data points are shown as individual dots, with outliers shown above or below the whiskers. P-values are indicated above statistically significant datasets.

### Chemokine signaling regulates expression of PDGFB

To test the possibility that CXCL12/CXCR4 signaling acts indirectly on vSMC motility via autocrine up-regulation of PDGFB in arterial endothelium (Fig. 4G), we examined whether chemokine signaling regulates PDGFB transcript and/or protein levels in HUVECs *in vitro* (Fig. 5A-E). Application of exogenous recombinant CXCL12 ligand to HUVECs in culture results in increased levels of PDGFB transcript (Fig. 5A) and PDGFB protein (Fig. 5B). Conversely, siRNA knockdown of endogenous CXCR4 or CXCL12 in cultured HUVECs results in decreased PDGFB transcript levels (Fig. 5C) and decreased PDGFB protein levels (Fig. 5D,E). To determine whether Cxcr4 is also required for Pdgfb expression in the endothelium *in vivo,* we used immunostaining to examine Pdgfb expression in the developing arteries of Cxcr4 knockout mice. Compared to their heterozygous siblings, Cxcr4-/- homozygous null mice showed a strong reduction in arterial expression of Pdgfb ligand, as well as reduced vSMC coverage as assessed by Sm22+ staining at E12.5 (Fig. 5F,G). Conversely, overexpression of *cxcl12b* ligand in zebrafish larvae via injection of *cxcl12b* RNA resulted in an increase in *pdgfb* transcript (Fig. 5H) and protein levels (Fig. 5I) as assessed by *in situ* hybridization and western blotting respectively. Together, these results suggest that autocrine CXCL12/CXCR4 chemokine signaling up-regulates PDGFB ligand in endothelial cells to promote recruitment of vSMC (Fig. 5J).

**Figure 5.**
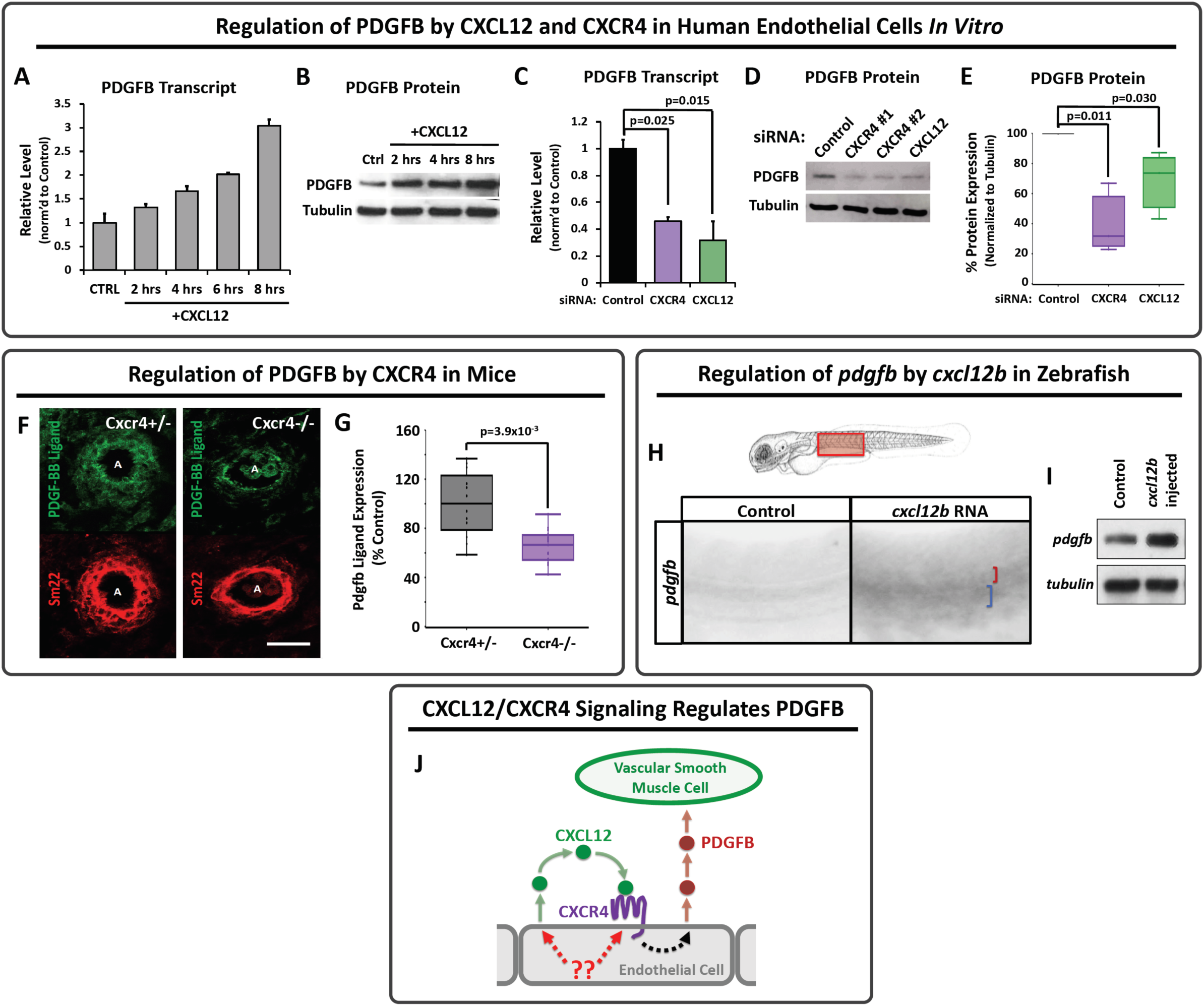
Chemokine signaling regulates PDGFB transcript and protein levels across species. **A,B,** PDGFB transcript (A) and protein (B) in HUVEC cells cultured *in vitro* in a confluent cell monolayer for up to 8 hours with (“+CXCL12”) or without (“CTRL”) added recombinant CXCL12. Relative PDGFB transcript levels (A) and protein levels (B) were measured by qPCR and Western blot, respectively, showing an upregulation of both PDGFB transcript and PDGFB protein levels in response to stimulation by CXCL12. **C-E**, PDGFB transcript (C) and protein levels (D,E) in HUVEC cells cultured *in vitro* in a confluent cell monolayer and treated with either control, CXCR4, or CXCL12 siRNAs. Relative PDGFB transcript (C) and protein (E,F) levels were measured by qPCR and Western blot, respectively, showing suppression of both PDGFB transcript and protein in response to either CXCR4 or CXCL12 knockdown. Values in A, C, and E are averaged from three individual experiments and expressed as a percentage of control. Error bars ± s.d. (A,C). **F,** Confocal images of immunohistochemically stained transverse sections through the dorsal aorta of E12.5 Cxcr4+/- heterozygous sibling (F) and Cxcr4-/- mutant (G) mice, probed for platelet derived growth factor B (PDGFB; green) and for smooth muscle 22 alpha (SM22, aka transgelin) for vascular smooth muscle cells (vSMC, red). **G,** Quantification of relative PDGFB protein expression in Cxcr4+/- heterozygous embryos versus Cxcr4-/- homozygous mutant embryos. Values are expressed as a percentage of heterozygous control and averaged from five individual mice per condition. **H**, Schematic diagram of a zebrafish larva with the red box highlighting the area imaged in panels I and J. **I,J,** Whole mount *in situ* hybridization of the mid-trunk of 2.5 dpf zebrafish injected with control (I) or *cxcl12b* (J) RNA, showing upregulation of *pdgfb* transcript in response to exogenous *cxcl12b*. Red and blue brackets in panel J indicate the dorsal aorta and cardinal vein, respectively. **K,** Western blot of whole embryo protein lysate from 2.5 dpf zebrafish injected with either control (left) or *cxcl12b* (right) RNA, probed for *pdgfb* (top) or alpha tubulin (bottom), showing upregulation of *pdgfb* protein levels in response to exogenous *cxcl12b*. Images are representative of data from three individual experiments. **M**, Schematic diagram illustrating the proposed model for endothelial-autonomous chemokine signaling driving increased endothelial PDGFB ligand production, thereby indirectly promoting vSMC acquisition by arteries. Putative upstream regulators of CXCL12 and CXCR4 are noted in red. Scale bars = 50 µm (panel F). Box plots are graphed showing the median versus the first and third quartiles of the data (the middle, top, and bottom lines of the box respectively). The whiskers demonstrate the spread of data within 1.5x above and below the interquartile range. All data points are shown as individual dots, with outliers shown above or below the whiskers. P-values are indicated above statistically significant datasets.

### KLF2 regulates vSMC recruitment upstream from chemokine signaling

We next sought to understand what might be responsible for preferentially restricting CXCL12/CXCR4 signaling to the dorsal aorta rather than the cardinal vein. Our primary candidate gene of interest was Klf2—a well documented blood flow-regulated transcription factor, with reported links to chemokine signaling. In the zebrafish, *klf2a* begins to be expressed in the cardinal vein just after the onset of blood flow between 24 and 32 hpf, and it remains expressed predominantly by veins through at least 72 hpf (Fig. 6A). Preferential expression of Klf2 in primitive veins is also noted in E9.5 mice (Fig. 6B), with Klf2 expression enriched in both the cardinal vein and the vitelline vein as compared to the dorsal aorta. These results, and previous reports that KLF2 can negatively regulate chemokine signaling, suggested to us that KLF2 expression in primitive veins might act as a “STOP” signal to block CXCL12/CXCR4 and PDGFB expression in the cardinal vein, thus restricting vSMC recruitment to the dorsal aorta. If this were the case, we predicted removing KLF2 would increase chemokine signaling and PDGFB expression to promote ectopic venous recruitment of vSMC.

**Figure 6.**
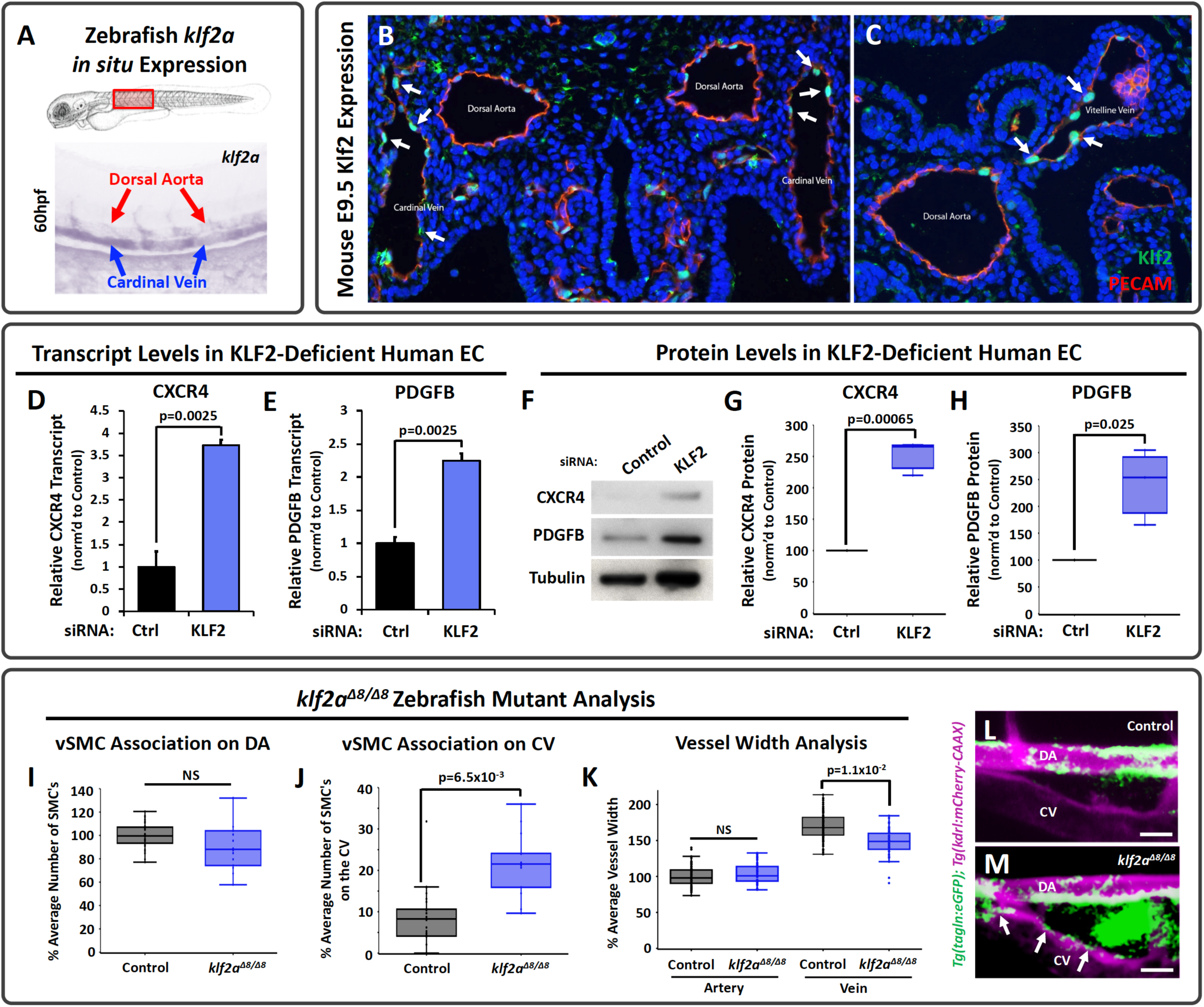
KLF2 is a negative regulator of chemokine signaling and PDGFB during early development. **A**) Top: Schematic diagram of a zebrafish larva with the red box highlighting the area imaged below. Bottom: Representative whole mount *in situ* hybridization (WISH) image of a 60 hpf zebrafish probed for *klf2a*. Red arrows indicate the dorsal aorta; blue arrows indicate the cardinal vein. **B,C**, Fluorescent images were taken of transverse sections through E9.5 mice: IHC for GFP was done to amplify signal from a Klf2-GFP knockin allele where the GFP is fused to the N-terminus of Klf2 (Klf2; green) and for platelet endothelial cell adhesion molecule-1 (PECAM) to mark the endothelium (red). Nuclei are labeled with DAPI (blue). Arrows highlight Klf2 positive endothelial nuclei in the cardinal vein (B) and vitelline vein (C) (Goddard et al., 2017; Weinreich et al., 2009). **D,E**, Quantitative qPCR measurement of CXCR4 (D) and PDGFB (E) transcript levels in HUVEC cells cultured *in vitro* as a confluent cell monolayer and treated with either control (black columns) or KLF2 (blue columns) siRNA. Values are expressed as a percentage of control. Error bars ± s.d. **F-H**, Representative Western blot images of CXCR4 and PDGFB protein levels (F), and quantification of relative CXCR4 (G) and PDGFB protein levels (H) from HUVEC cells cultured *in vitro* in a confluent cell monolayer and treated with either control or KLF2 siRNA. Values in G,H are averaged from three individual experiments and expressed as a percentage of control. **I,J,** Quantification of vSMC number associated with the dorsal aorta (I) or cardinal vein (J) in the mid-trunk of 5 dpf wild type sibling (black columns) or *klf2a^Δ8/Δ8^* mutant (blue columns) animals. Values are averaged from three individual experiments and expressed as a percentage of wild type siblings. **K**, Quantification of dorsal aorta (DA, left columns) and cardinal vein (CV, right columns) width in the mid-trunk of 5 dpf wild type sibling (black columns) or *klf2a^Δ8/Δ8^* mutant (blue columns) animals. **L,M**, Confocal images of the anterior trunk of 5 dpf *Tg(tagln:eGFP), Tg(kdrl:mCherry-CAAX)* sibling (L) and *klf2a^Δ8/Δ8^* mutant (M) zebrafish embryos with eGFP positive vSMCs (green) and mCherry-CAAX positive endothelial cells (magenta). Arrows point to vSMCs associated with the CV in *klf2a^Δ8/Δ8^* mutants. Scale bars = 75 µm (panels L,M). Box plots are graphed showing the median versus the first and third quartiles of the data (the middle, top, and bottom lines of the box respectively). The whiskers demonstrate the spread of data within 1.5x above and below the interquartile range. All data points are shown as individual dots, with outliers shown above or below the whiskers. P-values are indicated above statistically significant datasets.

To test this idea, we used siRNA knockdown to suppress KLF2 in HUVECs *in vitro* and genetic mutants to “knock out” *klf2a* in the zebrafish *in vivo.* Suppressing KLF2 in HUVECs *in vitro* led to upregulation of CXCR4 and PDGFB transcript and protein levels (Fig. 6D-H). To examine the consequences of reduced KLF2 in the endothelium *in vivo*, we used CRISPR/Cas9 mutagenesis to generate an 8 bp deletion mutant in zebrafish *klf2a* (Supp. Fig. 1D). Although there was no change in the number of vSMCs associated with the dorsal aorta in homozygous *klf2a^Δ8/Δ8^* mutants (Fig. 6I,L,M), there was a clear increase in the number of vSMCs associated with the cardinal vein (Fig. 6J,L,M). This increase was accompanied by a modest decrease in the diameter of the cardinal vein without any change in the dorsal aorta (Fig. 6K), consistent with the role of vSMC in regulating vessel diameter. Together, these results suggest that expression of KLF2 prevents chemokine/PDGF-mediated association of vSMC to primitive veins.

## DISCUSSION

### A molecular pathway for vSMC recruitment

The preferential association of vascular smooth muscle cells with arteries has been appreciated for centuries, but the molecular mechanisms underlying this preference have not been fully elucidated. We have uncovered a molecular pathway that promotes association of vSMCs with developing arteries and reduces vSMC association with veins (Fig. 7). The known blood flow-regulated transcription factor KLF2 helps demarcate and maintain a pro-vSMC state in the endothelium. Sustained expression of KLF2 (zebrafish *klf2a*) in primitive venous endothelium during early development represses expression of *cxcl12b* and *cxcr4a* and autocrine/juxtacrine endothelial chemokine signaling, thereby reducing chemokine signaling-promoted production of *pdgfb* and venous recruitment of vSMC. In contrast, down-regulation of *klf2a* in the early arterial endothelium permits expression of *cxcl12b* and *cxcr4a*, promoting production of arterial *pdgfb* and arterial recruitment of vSMC (Fig. 7). As predicted by this model, either artificially suppressing *klf2a* expression or mis-expressing *cxcl12b* in venous endothelium promotes ectopic venous recruitment of vSMC (Fig. 3A-E, Fig. 6J).

**Figure 7.**
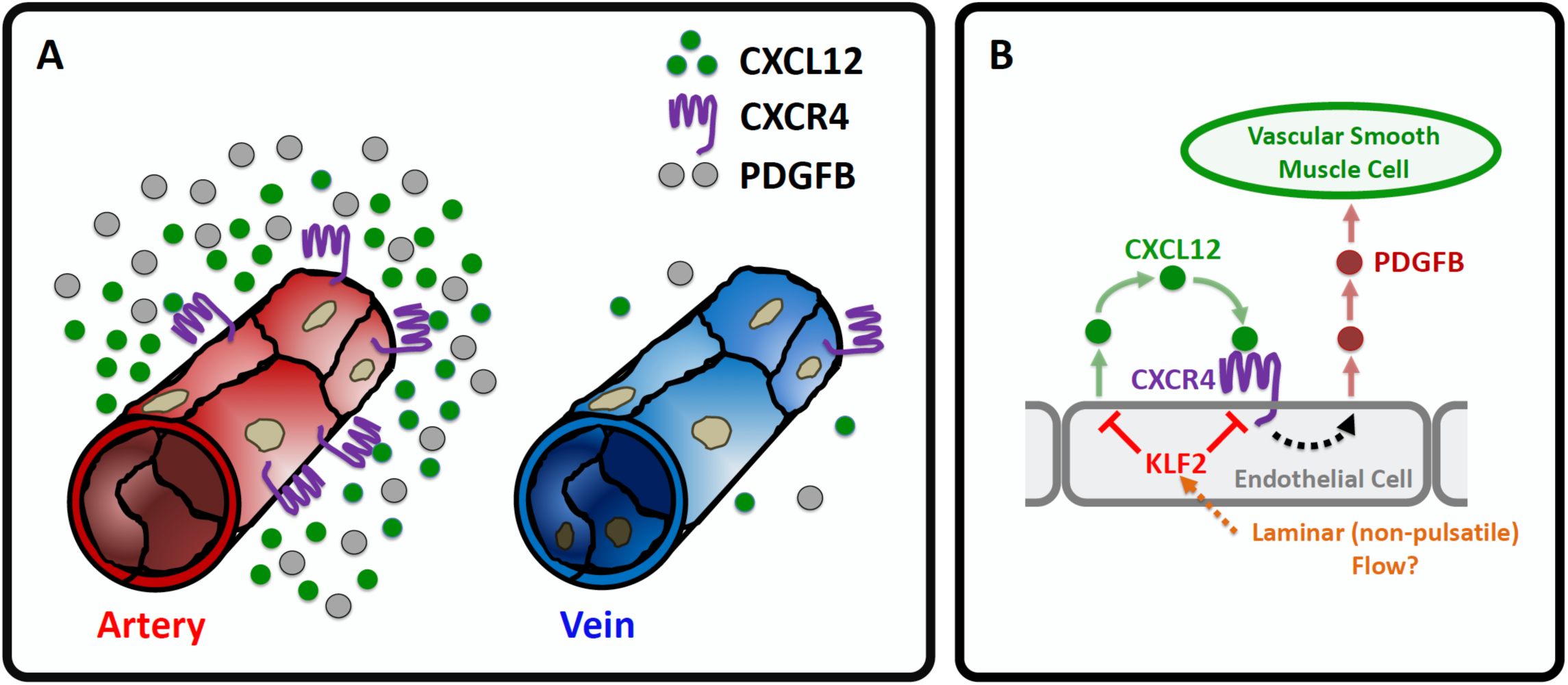
A proposed model for the preferential recruitment of vascular smooth muscle cells to arteries. **A**, During the early development of arterial and venous blood vessels the CXCL12 ligand, its receptor CXCR4, and the vascular smooth muscle cell (vSMC) chemoattractant PDGFB are all more highly expressed on arteries than on veins. In contrast, KLF2 is more highly expressed on primitive veins than arteries. **B,** A proposed molecular pathway for preferential recruitment of vSMC to arteries. In arteries, autocrine activation of endothelial CXCR4 by its CXCL12 ligand results in increased production of PDGFB by arterial endothelium, promoting vSMC recruitment to arteries. Expression of KLF2 in primitive veins suppresses expression of CXCL12 and CXCR4, preventing up-regulation of PDGFB and limiting vSMC recruitment to veins. The cues directing preferential expression of KLF2 in veins remain unclear, but may involve different types of flow (ie, pulsatile vs. laminar).

### The role of blood flow in vSMC recruitment and differentiation

Together, our findings and previous reports describing flow-mediated regulation of KLF2 suggest that changes in blood flow velocity, force, or directionality influence stabilization of the arterial blood vessel wall by modulating the expression of this key transcription factor. In the zebrafish trunk, *klf2a* expression is initially equivalently low in the endothelial cords that will give rise to the dorsal aorta (DA) and cardinal vein (CV), but as these vessels lumenize and circulation begins, the expression of *klf2a* is strengthened in the CV. Since flow begins in the DA and CV at the same time during early development, why do these two vessels show such different *klf2a* expression, and how could flow be playing a role in this difference? Although further work will be needed to definitively answer this question, the *type* of flow that each of these vessels experience is likely playing a critical role in their differential expression of *klf2a*. There is relatively even, constant laminar flow through the CV at these early developmental stages, while the DA experiences a much more pulsatile flow regime. Pulsatile flow results in cyclical changes in velocity and shear, and in the circumferential “stretch” experienced by the endothelial cells in the DA, that are not experienced by CV endothelial cells. Indeed, our previously published findings show a high degree of pulsatility and cyclical change in diameter of the DA with blood flow pulsation during early development (Stratman et al., 2017). *In vitro* and *in vivo* studies have shown that KLF2 expression is differentially regulated by different types of flow regimes—typically showing increased expression by laminar flow and reduced expression associated turbulent, non-laminar flow (Goddard et al., 2017; Lee et al., 2006; Lin et al., 2010; Methe et al., 2007; Nayak et al., 2011; Parmar et al., 2006).

Although we show that developing primitive arteries show reduced KLF2 expression compared to primitive veins in both zebrafish and mice, the endothelium of mature arteries expresses high levels of KLF2 (Goddard et al., 2017; Methe et al., 2007; Nayak et al., 2011; Parmar et al., 2006). How can the latter results be reconciled with our model (Fig. 7) in which “arterial” blood flow reduces KLF2 expression and the presence of KLF2 suppresses arterial-specific recruitment of vSMC? These results make sense if the expression of KLF2 is responsive to endothelial cell stretch/strain. Primitive arteries initially only consist of an endothelial layer without encircling smooth muscle layers, resulting in high levels of endothelial stretch and strain in response to pulsatile flow, as noted above. Gradual acquisition of vSMCs and assembly of vascular basement membrane layers creates resistance to the cyclical changes in vessel diameter with blood flow pulsation, reducing the “stretch” experienced by endothelial cells. Again, our previously published results show reduced vessel wall movements and pulsatile blood flow-induced changes in diameter as the DA acquires vSMC (Stratman et al., 2017). Endothelial recruitment of vSMC is required for formation of a vascular basement membrane, and this is also critical for reducing vessel distensibility, and thus endothelial cell stretch/strain (Stratman and Davis, 2012; Stratman et al., 2009; Stratman et al., 2017; Stratman et al., 2010).

Together, these data point to dynamic, flow-regulated expression of KLF2 as a mechanism not only for promoting arterial-specific recruitment of vSMC, but also for self-limiting and self-tuning the extent of vSMC acquisition to match the specific hemodynamic requirements of different regions of the arterial vascular tree. As developing arteries acquire vSMC and vSMC-promoted basement membrane, reduced vessel compliance/increased stiffness would be predicted to result in increased KLF2 expression, creating a “stop signal” to halt further vSMC recruitment and promote vSMC differentiation and quiescence (Adam et al., 2000; Liu et al., 2005; Sebzda et al., 2008). In larger and/or more proximal arteries experiencing more dramatic cyclic flux in blood pressure, the “stop signal” might be expected to come later, after acquisition of a greater amount of vSMC and matrix and formation of a thicker and more robust vessel wall. Thus, this mechanism would ensure that the capacity of the vascular wall to resist stretch matches the flow regime vessels experience. Although this seems like a very attractive mechanism to allow arteries to “self-regulate” vSMC acquisition based on intrinsic flow dynamics, additional experimental studies will of course be needed to further substantiate that this is indeed an important determinant of the extent of vascular wall assembly.

### Developmental pathways reactivated in disease?

A number of disease states have known links to altered blood flow sensing and/or are correlated with altered Klf2 expression profiles, including atherosclerosis and Cerebral Cavernous Malformations (CCMs), and it seems possible that the KLF2-CXCL12/CXCR4-PDGFB molecular pathway that we have identified for arterial vSMC acquisition during development may also play critical roles in vascular pathologies. As noted in our introduction, sites of low Klf2 expression are correlated with atherosclerotic lesion formation (Atkins and Jain, 2007; Komaravolu et al., 2015; Lee et al., 2006; Parmar et al., 2006). The experimental data from our study showing that low Klf2 expression correlates with high chemokine and high PDGFB expression suggests that suppressed Klf2 could be promoting atherogenesis at least in part by promoting chemokine/PDGFB-stimulated vSMCs proliferation and reactivated motility (Adam et al., 2000; Dandre and Owens, 2004; Doring et al., 2014; Liu et al., 2005; Noels et al., 2014; Owens et al., 2004; Sebzda et al., 2008; Sen-Banerjee et al., 2005). Altered chemokine signaling has already been implicated in coronary artery disease, mobilization of progenitor cells in ischemia induced remodeling/repair, and atherosclerosis (Busillo and Benovic, 2007; Doring et al., 2014; Gupta et al., 1998; Noels et al., 2014; Sebzda et al., 2008; Shi et al., 2014). Upregulated chemokine signaling can lead to increased smooth muscle cell proliferation and chemotaxis, and it is known that vSMCs actively de-differentiate and invade atherosclerotic plaques (Doring et al., 2014; Noels et al., 2014). Our results would suggest that the effects of chemokine signaling on the motility and proliferation of vSMCs in these pathological contexts may be indirect, through upregulation of PDGFB. Other disease models also offer some support for this idea.

CCM pathology, on the other hand, is associated with high Klf2/4 expression (Li et al., 2019; Renz et al., 2015; Zhou et al., 2016). CCM is a vascular malformation disorder particularly localized to intersection points between capillaries and veins that leads to the formation of large dilated vessels (Castro et al., 2019; Li et al., 2019; Renz et al., 2015; Zhou et al., 2016). These dilated vessels are prone to rupture, leading to trauma, tissue damage, and ischemia. Formation of CCM lesions has been linked to mutations in one of three different CCM proteins, that appear to have their deleterious effects at least in part via a Cdc42-MEKK3-MEK5-ERK5-KLF2/4 signaling cascade within the endothelium (Castro et al., 2019; Komaravolu et al., 2015; Li et al., 2019; Parmar et al., 2006; Zhou et al., 2016). Vessels in CCM lesions have low numbers of associated vSMC/mural cells, consistent with the idea that high Klf2 levels serve as a “stop” signal for vSMC recruitment. It will be interesting to further determine whether vSMC recruitment and the KLF2-CXCL12/CXCR4-PDGFB pathway we have identified are involved in a causative way in lesion formation, or only secondarily affected as a result and/or part of developing CCM lesion pathology.

### Concluding Remarks

We describe a signaling pathway that promotes association of vSMCs with the arterial vasculature and restrains association with the venous vasculature during development. This pathway links blood flow, through the transcription factor Klf2, to chemokine signaling, PDGFB production, and ultimately arterial endothelium-directed motility of vSMCs. In addition to its important role in arterial/venous development, this signaling cascade may also be involved in vascular pathologies that alter vSMC acquisition or function.

## METHODS

### Zebrafish Methods and Zebrafish Transgenic Lines

Zebrafish *(Danio rerio)* embryos were raised and maintained as described (Kimmel et al., 1995; Westerfield, 1995). Zebrafish husbandry and research protocols were reviewed and approved by the NICHD Animal Care and Use Committee. Zebrafish transgenic lines Tg*(kdrl:mCherry-CAAX) y171*; Tg(*tagln:egfp*)*p151* are previously published. New CRISPR/Cas9 mutant alleles generated for this manuscript include: (*cxcl12b^Δ24^)y611*; *(cxcr4a^Δ7^)y612*; *(pdgfbb^Δ3^(chromosome3))y613*; *(pdgfba^Δ6^ (chromosome22))y614*; *(klf2a^Δ8^)y616*

### Reagents

Antibodies for immunostaining and western blot analysis include: Tubulin (Sigma, #T6199-1:10,000 dilution); PECAM-1 (CD31) (BD Pharmingen; #553370-1:300 dilution); alpha-sma-cy3 (Sigma; #c-6198; 1:500 dilution); Sm22 (GeneTex; #GTX101608; 1:300 dilution); PDGFB (SantaCruz; # sc-365805; 1:500 dilution); CXCR4 (Sigma, #SAB3500383); CXCL12 (R&D Systems, #AF-310-NA). WISH probes utilized include *klf2a* (Corti et al., 2011), *pdgfb* (Wiens et al., 2010), *cxcr4a* (Cha et al., 2012; Fujita et al., 2011), and *cxcl12b* (Primers FW:GAGCTCT GGACACTCGCTGT; RV:TACTGCTGAAGCCATTTGGTC)

### Imaging and Microscopy

Fluorescent images were collected utilizing either the Leica SP5 II or Nikon Yokogawa CSU-W1 spinning disk confocal microscope at 5dpf at 20x magnification. Embryos were immobilized in buffered MS-222 and embedded in 0.8% low melting point agarose. All images were acquired, data quantified and analyzed blindly, and then embryos genotyped.

### Endothelial Cell Culture and 3D Assays

Human umbilical vein endothelial cells (HUVEC, Lonza) were cultured in bovine hypothalamus extract, 0.01% Heparin and 20% FBS in M199 base media (Gibco) on 1mg/ml gelatin coated tissue culture flasks. HUVECs were used from passages 3-6.

Human coronary artery smooth muscle cells (PASMC, Lonza) were cultured in 10% FBS in Advanced DMEM base media (Gibco) on 1mg/ml gelatin coated tissue culture flasks. PASMCs were used from passages 3-8.

3-dimensional (3D) collagen type I *in vitro* assays were done essentially as described ((Stratman et al., 2009)), utilizing 2.5mg/mL collagen type I (BD Biosciences, Acid Extracted) gels including CXCL12 (R&D Systems, #350-NS/CF) or PDGFB (R&D Systems, #220-BB/CF). PASMCs were seeded on the collagen gel at 40,000 cells per well density. Culture media for the assays contained ascorbic acid, FGF (R&D Systems, #233-FB-025/CF), and IGF-II (R&D Systems, #292-G2-250). Assays were fixed in 2% paraformaldehyde (PFA) at 2 days and processed for future analysis.

### CRISPR/Cas9 Generation of Zebrafish Mutants

Mutations in the zebrafish *cxcl12b, cxcr4a, pdgfbb, pdgfba,* and *klf2a* genes were generated using the CRISPR/Cas9 system. The following guide RNAs were transcribed *in vitro* using the T7 mMessage Machine® Kit (Ambion), and injected at a dose of 150 pg/nl per embryo:

cxcl12b^Δ24/Δ24^:

TAATACGACTCACTATAGGAGCCCAGAGACTGACGGTGTTTTAGAGCTAGAA

*cxcr4a^Δ7/Δ7^*:

TAATACGACTCACTATAGGACATCGGAGCCAACTTTGGTTTTAGAGCTAGAAATAGC

AAG

*pdgfbb*:

TAATACGACTCACTATAGGCTGTGGTTGAGTTGGTGAGTTTTAGAGCTAGAA

*pdgfba*:

TAATACGACTCACTATAGGACCCTCTTCCTCCATCTCGTTTTAGAGCTAGAA

*klf2a^Δ8/Δ8^*:

TAATACGACTCACTATAGGTCCGTAACTATCCATGCAGTTTTAGAGCTAGAAATAGC

AAG

pT3TS-nCas9 (Addgene) was transcribed using MEGA Script T7 kit (Invitrogen/Ambion), and injected at a dose of 300pg/nl per embryo. Embryos were injected at the single cell stage, screened for cutting efficiency and grown on system. F1 generations were analyzed for mutations, and pairs crossed for analysis in the F2 and beyond generations.

### Genotyping of Zebrafish Mutants

Mutants were genotyped using the following primers:

*cxcl12b^Δ24^*:

FW: TGTAAAACGACGGCCAGTGTATCACTTATATTCTCAAC

RV: GTGTCTTCACTCGCTCTTGGCATGGATAGC

*cxcr4a^Δ7^*:

FW: TGTAAAACGACGGCCAGTCAGCACATCGTCTTTGAAGATGATTTATC

RV: GTGTCTTGGCAGAGTGAGCACAAACAGAAGG

*pdgfb ^Δ4^ (chromosome 3)*:

FW: TGTAAAACGACGGCCAGTGATTGTTTGATTAATAAGGAC

RV: GTGTCTTCTACAACATGTGACAAATTC

*pdgfa ^Δ8^ or ^Δ20^ (chromosome 22)*:

FW: TGTAAAACGACGGCCAGTAGGTGTTGTTTTGTTCAGGACC

RV: GTGTCTTTGGTATGGGATCAGCTTTACCT

*klf2a^Δ8^*:

FW: TGTAAAACGACGGCCAGTGACATTGACACCTACTGC

RV: GTGTCTTGAGTCATGCTGCCTGCTCC

Universal Primer: FAM-M13: 5’-/56-FAM/ TGTAAAACGACGGCCAGT -3’

### ABI 3130xl Fragment Analyzer Protocol

PCR protocol with AmpliTaq Gold DNA Polymerase 1x (10ul) Rxn: 1ul 10x PCR Gold Buffer; 0.5ul MgCl2 25mM; 1ul 0.5mM ABI Fwd primer; 1ul 1mM ABI Rev primer; 0.2ul 10mM FAM-M13 primer; 0.1ul dNTP Master Mix; 0.1ul TaqGold polymerase; 1ul of 1:10 diluted crude gDNA; 5.1ul H20

TaqGold PCR Program: 95oC 10min’ 95oC 30sec; 58oC 30sec; 72oC 30sec (1 min/kb); GoTo Step 2 x34; 72oC 10min; 15oC Hold; Run on ABI immediately or store at 4oC in the dark for 24 hours max.

ABI 3130xl Plate set-up: HiDi Formamide/ROX master mix-0.2ul ROX400HD; 9.8ul HiDi Formamide; add 10ul of master mix to each ABI plate sample well; add 2ul of fluorescent PCR product; cap wells and denature at 95oC for 5min; uncap all wells and replace with ABI plate septa to run on the 3130xl. Follow manufacturer directions to utilize the ABI 3130xl.

### Mouse Lines, Breeding, and Genotyping

Cxcr4 Knockout Mice: *Cxcr4* knockout mutants: We initially generated *Cxcr4+/-* heterozygous mice from the breeding of *Cxcr4-flox* mice (Jax# 008767) with *E2a-Cre* mice (Jax#003724). *Cxcr4-/-* homozygous embryos were produced by mating *Cxcr4+/-* heterozygous mice. Offspring were genotyped by genomic PCR using primers that specifically detect *Cxcr4* allele and *Cxcr4* null allele.

*Cxcr4* allele (347bp):

FW: CAC TAC GCA TGA CTC GAA ATG

RV: GTG TGC GGT GGT ATC CAG C

*Cxcr4* null allele (190bp)

FW: CAC TAC GCA TGA CTC GAA ATG

RV: CCT CGG AAT GAA GAG ATT ATG C

Klf2-GFP mouse was published previously (Weinreich et al., 2009), and is a knockin allele where the GFP is fused to the N-terminus of KLF2. GFP signal was amplified using a GFP antibody (Goat anti-GFP, Abcam, CAT#ab6673 1:250) and the endothelium labeled with PECAM-1 antibody (Dianova, CAT# DIA-310 1:200).

### Immunostaining and Western blot analysis

Tissue sections were immunostained following the same basic protocol: 1) 30 min RT incubation in Tris-Glycine; 2) 1hr RT incubation +/- permeabilization with 0.01% TritonX-100; 3) 2hr RT incubation in blocking solution (5% Sheep Serum, 1% Roche Blocking Buffer in PBST); 4) 1hr at RT - overnight 4oC incubation with 1:1000 primary antibody unless otherwise noted; 5) wash with PBST; 6) 2-3hr RT incubation with 1:2000 secondary antibody in 5% Sheep Serum, 1% Roche Blocking Buffer in PBST; 7) wash with PBST and imaging analysis. Quantification of immunostaining intensity was performed by ImageJ analysis software. Images were acquired using a Leica SP5 II confocal microscope or Nikon Yokogawa CSU-W1 spinning disk confocal microscope. All images were acquired at the same intensity, step size and image resolution for analysis. The images were analyzed using ImageJ (Schindelin et al., 2012). Data is reported as the percent average intensity per region of interest size from a minimum of 3 images from 3 independent experiments/animals ± s.e.m.

Zebrafish samples for Western blot analysis were deyolked and directly lysed in 2x Laemmli Sample Buffer containing 5% b-ME and a PhosSTOP tablet (Roche), 10 ul per embryo unless otherwise indicated. HUVEC cells for Western blot analysis were lysed directly in 2x Laemmli Sample Buffer containing 5% b-ME and a PhosSTOP tablet (Roche), 500ul per T-25 culture flask.

Secondary Digital-HRP-conjugated antibodies were purchased from Kindle Bioscience and used at 1:1000 in 5% milk. Primary antibodies are described in the ‘Reagents’ section and used at 1:1000 unless otherwise noted. Images were acquired using the Kindle Bioscience KwikQuant Imager and 1-Shot Digital ECL. Quantification of relative band density was performed using ImageJ software. Data is reported as the percent average density from a minimum of 2-3 blots from at least 2 independent experiments ± s.e.m.

### qPCR and RNA Extraction

Zebrafish embryos or HUVECs were collected at the indicated time points in TRIZOL and RNA purified using a double chloroform extraction protocol. cDNA was generated using BioRads I-Script cDNA synthesis kit from 500ng of RNA. TaqMan qPCR protocols were utilized to generate relative expression data, and analysis run using the FAM channel of a 96 well BioRad CFX qPCR machine. Primer product numbers are as follow:

PDGFB (human): Hs00966522_m1

CXCL12 (human): Hs03676656_mH

KLF2 (human): Hs00360439_g1

CXCR4 (human): Hs00607978_s1

Ef1a (human): Hs00265885_g1

GAPDH (human): Hs02758991_g1

### Statistics

Statistical analysis of data was done using Microsoft Excel. Statistical significance was set at a minimum of p ≤ 0.05 and is indicated in individual figures. Student’s t-tests were used when analyzing 2 groups within a single experiment. Bar graphs were generated with Plotly.

### Study Approval

Zebrafish husbandry and research protocols were reviewed and approved by the NICHD Animal Care and Use Committee at the National Institutes of Health. All animal studies were carried out according to NIH-approved protocols, in compliance with the *Guide for the Care and use of Laboratory Animals*.

## Author Contributions

ANS, OMF, MCB, AED, WL, VNP, DC, ON, JY, TJB, LG, and MVG performed experiments; ANS, OMF, MCB, AED, WL, VNP, DC, ON, JY, LG, MVG, TJB, MK, YSM, and BMW analyzed results and made the figures; ANS, OMF, MCB, AED, WL, VNP, DC, JY, MK, YSM, and BMW designed the research and wrote the paper.

## Correspondence

Brant M. Weinstein, Department of Developmental Biology, National Institute of Child Health and Human Development, National Institutes of Health, 6 Center Dr. Bethesda, MD 20892;

## Figure Legends

**Supplemental Figure 1.**
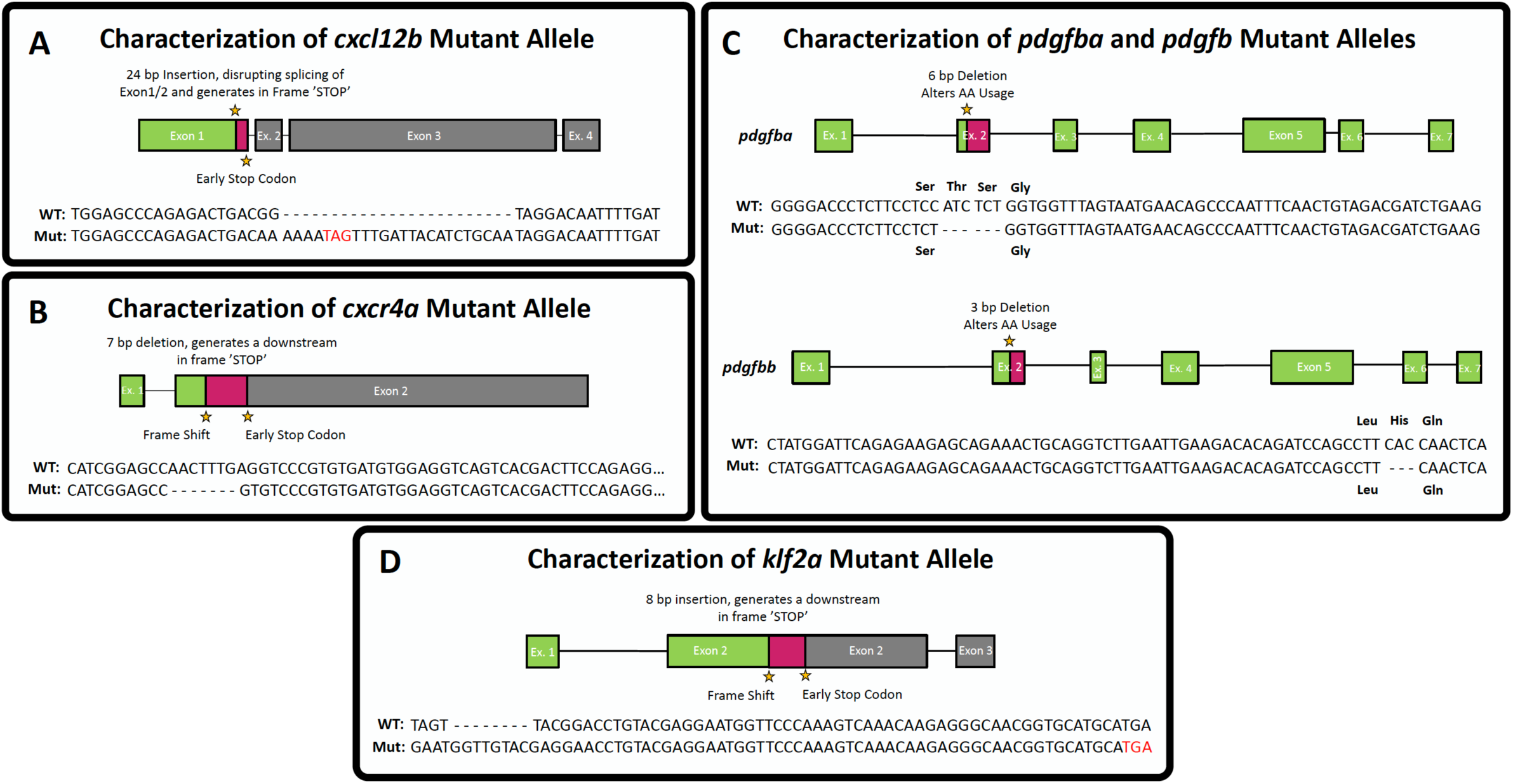
CRISPR/Cas9 mutant alleles. Gene structure and sequence of mutants generated for this study using CRISPR/Cas9 mutagenesis. **A, *cxcl12b^Δ24^* mutant.** A 24 base pair insertion was introduced that destroys the splice donor site between exons 1 and 2 and inserts an in frame early stop codon. **B, *cxcr4^Δ7^* mutant.** A 7 base pair deletion in exon 2 leads to a frame shift at amino acid 75 and early termination after amino acid 198. **C, *pdgfba ^Δ6^* mutant.** A 6 base pair deletion in exon 2 (chr22:29,325,360-29,325,754) leads to an in-frame deletion of two polar amino acids-a threonine and serine-leading to predicted alterations in protein folding. **D, *pdgfbb^Δ3^* mutant.** A 3 base pair deletion in exon 2 that leads to an in-frame deletion of a histidine leading to predicted alterations in protein folding. **E, *klf2a^Δ8^* mutant.** An 8 base pair insertion in exon 2 leads to a frame shift at amino acid 476 and early termination after amino acid 543.

**Supplemental Figure 2.**
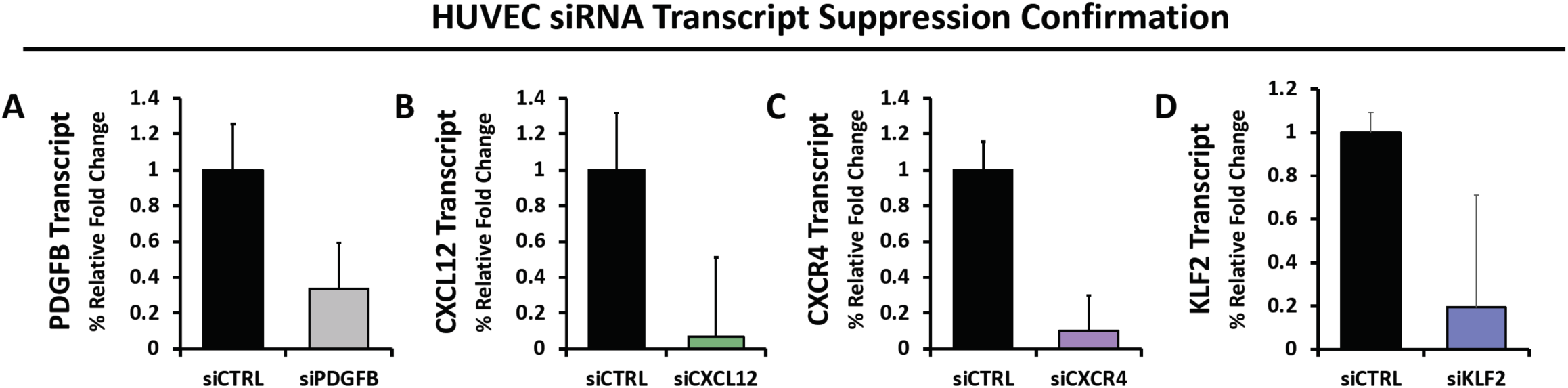
siRNA knockdown in Human Umbilical Vein Endothelial Cells (HUVEC). We confirmed suppression of genes targeted by siRNAs in HUVECs for this study. Transcript levels were measured by performing qRT-PCR for each respective gene, comparing control siRNA treated to PDGFB (**A**), CXCL12 (**B**), CXCR4 (**C**), or KLF2 (**D**) siRNA treated cell samples. Data are representative of 3 experimental replicates. Values are normalized to control siRNA levels. Error bars ± s.d.

## Acknowledgements

The authors would like to thank members of the Weinstein laboratory for their critical comments on this manuscript. This work was supported by the intramural program of the *Eunice Kennedy Shriver* National Institute of Child Health and Human Development, National Institutes of Health (ZIA-HD001011 and ZIA-HD008915, to BMW); K99/R00 Pathway to Independence Award (NHLBI-4R00HL125683-02, to ANS); the Intramural Research Program of the National Heart, Lung, and Blood Institute, National Institutes of Health (ZIA HL005702-14 to YM).

